# Targeting neurons in the tumor microenvironment with bupivacaine nanoparticles reduces breast cancer progression and metastases

**DOI:** 10.1101/2021.05.18.444598

**Authors:** Maya Kaduri, Mor Sela, Shaked Kagan, Maria Poley, Hanan Abumanhal-Masarweh, Patricia Mora-Raimundo, Alberto Ouro, Nitsan Dahan, Jeny Shklover, Janna Shainsky-Roitman, Yosef Buganim, Avi Schroeder

## Abstract

Neurons within the tumor microenvironment promote cancer progression, thus their local targeting has potential clinical benefits. We designed PEGylated lipid nanoparticles loaded with a non-opioid analgesic, bupivacaine, to target neurons within breast cancer tumors and suppress nerve-to-cancer crosstalk. *In vitro*, 100-nm nanoparticles were taken up readily by primary neurons, trafficking from the neuronal body and along the axons. We demonstrate that signaling between triple-negative breast cancer cells (4T1) and neurons involves secretion of cytokines stimulating neurite outgrowth. Reciprocally, neurons stimulated 4T1 proliferation, migration and survival through secretion of neurotransmitters. Bupivacaine curbs neurite growth and signaling with cancer cells, inhibiting cancer-cell viability. *In vivo*, bupivacaine-loaded nanoparticles administered intravenously, suppressed neurons in orthotopic triple-negative breast cancer tumors, inhibiting tumor growth and metastatic dissemination. Overall, our findings suggest that reducing nerve involvement in tumors is important for treating cancer.

## Introduction

Tumor-infiltrating nerves from the peripheral nervous system, occurs in various types of cancer. Nerve/cancer interactions have been shown to support cancer development and progression (*1*). Cancer cells secrete neurotrophic factors that promote nerve innervation into the tumor microenvironment in a process termed neurogenesis (*2*). Additionally, the nervous system has an essential impact on stimulating cancer-cell growth, proliferation, angiogenesis and invasion through the secretion of chemokines and neurotransmitters (*3*). Perineural invasion (PNI) (*4*) of peripheral nerves into tumors and the reciprocal interactions between cancer cells and nerves, suggest that targeting nerves in the tumor tissue will be beneficial for treating various cancer types, such as prostate, breast, lung, ovarian and pancreatic cancer (*1*). Chemical and surgical depletion of sympathetic nerves was shown to suppress the early stage of prostate cancer development (*5*). In addition, genetic downregulation of tumor infiltrating sympathetic nerves, suppressed tumor progression in breast cancer models (*6*). Moreover, β adrenergic blockers treatment reduced progression of breast, prostate, lung, ovarian and melanoma cancers (*7*). Therefore, various therapeutic approaches which manipulate the local nerves in the tumor tissue have inhibitory effect on cancer (*1, 8*).

Nanotechnology is gaining attention in targeted cancer therapies and diagnostics (*9-13*), as well as therapeutic applications in neuronal regeneration (*14*). Specifically, liposomes, self-assembled lipid vesicles, are commonly used as nanoscale drug delivery systems (*15, 16*). Bupivacaine is a non-opioid selective sodium channel blocker, that interrupts the transmission of the nerve impulse and pain signals. Encapsulation of bupivacaine into nanoparticles have been shown to reduce occurrence of systemic adverse effects from intravenously injection of bupivacaine (*17*). Micron-scale liposomes containing bupivacaine have been developed for prolonged local analgesia in the management of postsurgical pain (*17*). We hypothesized that bupivacaine loaded nanoparticles will suppress neuronal activity in the tumor microenvironment, to improve cancer management. We examined the nerve/cancer interactions and tested liposomal bupivacaine’s ability to curb nerve signaling in breast cancer *in vitro* and *in vivo* models.

## Results

The collaborative interactions between cancer and the nervous system promote cancer progression (*1*). In this study, lipid nanoparticles encapsulating the non-opioid analgesic, bupivacaine, were used to suppress these interactions, thus inhibiting tumor growth and metastasis in orthotopic triple negative breast cancer (4T1) murine models (Fig. 1A). *In vitro* models included cortical primary neurons and rat adrenal pheochromocytoma cells (PC12) (*18*).

**Figure 1.**
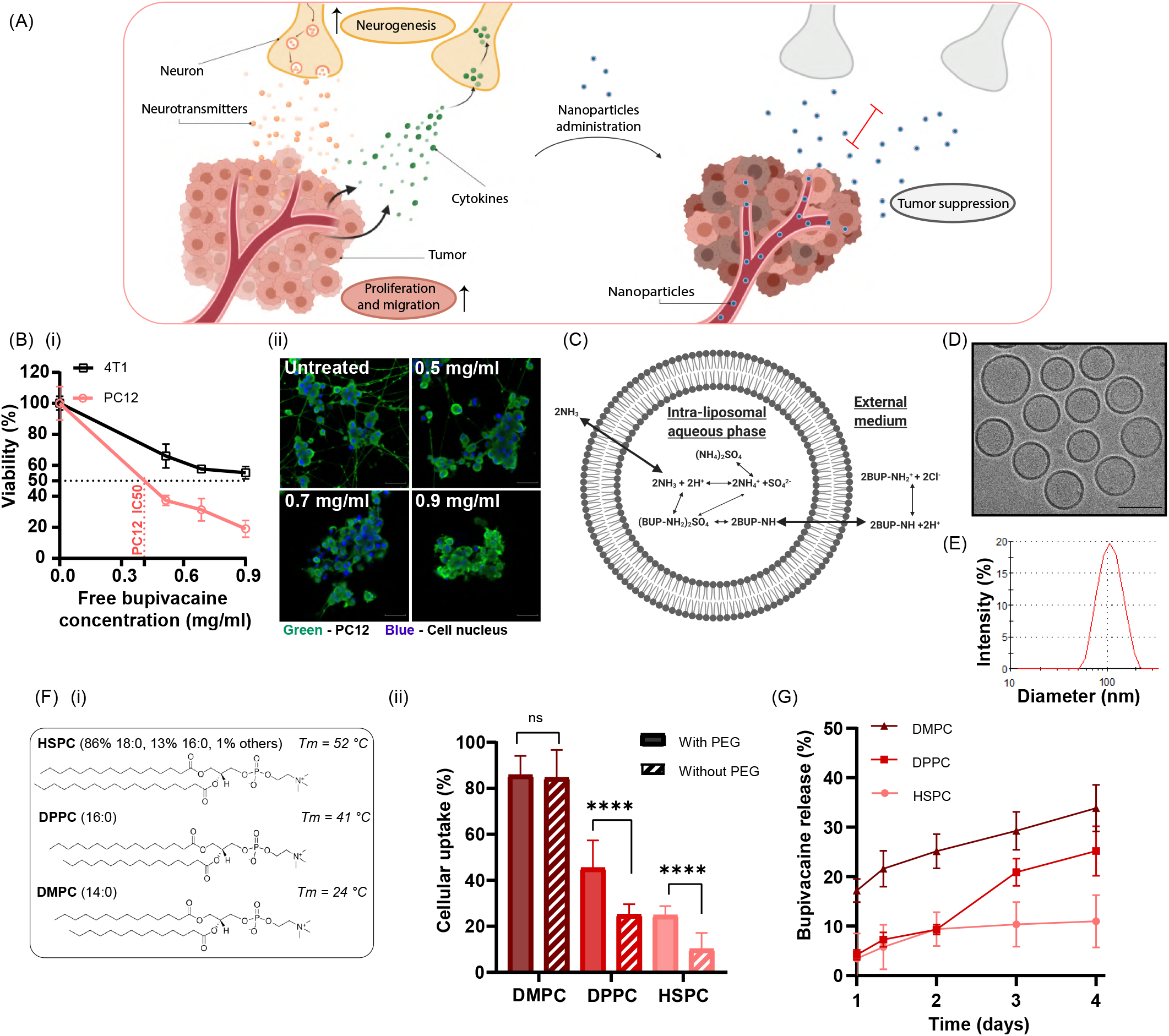
Analgesic nanoparticles as a tool for treating neurons within breast cancer tumors. Cross-talk between cancer cells and nerves supports cancer-cell proliferation and migration; reciprocally, neurite growth is promoted through secretion of cytokines by cancer cells. We used lipid-nanoparticles, liposomes, loaded with bupivacaine (L-BUP) to curb nerve/cancer cross-talk and inhibit tumor growth (**A**). Viability test of PC12 neurons and triple-negative breast cancer cells (4T1) after treatment with bupivacaine (**B-i**, normalized to the untreated group), and confocal imaging (**B-ii**, scale bar is 50µm). 100nm liposomes composed of HSPC, cholesterol and DSPE-PEG2000 (55:40:5 molar ratio) were loaded with bupivacaine using ammonium sulfate gradient (**C**), and characterized by cryo-TEM (**D**, scale bar is 100nm) and dynamic light scattering (**E**, PDI<0.1). Various lipid formulations were compared for their neuronal uptake: HSPC, DPPC, DMPC, with cholesterol and with/without DSPE-PEG2000 (**F, i**). PC12 cells were incubated with Rhodamine-labeled liposomes and analyzed for their cellular uptake (**F-ii**). Liposomal bupivacaine release profile was conducted at 37°C, comparing the stability of the different formulations (**G**). Results are presented as mean±SD (between 2 to 3 independent repetitions preformed in at least 3 replicates). Two-way ANOVA was used for statistical analysis of (B-i) and One-way ANOVA for (F-ii and G) with multiple comparisons test adjusted *p* value; ****p<0.0001.

### Neurotoxicity of bupivacaine

Bupivacaine, a selective sodium channel blocker, has a neurotoxic effect that is both concentration and duration dependent (*17*). The median lethal concentration (LC_50_) of bupivacaine was evaluated in cell lines for both, neurons (PC12 cells) and cancer (4T1 cells). For PC12 cells LC_50_ was ∼0.4 mg/ml compared to ∼0.9 mg/ml for 4T1, indicating a higher potency of the drug towards neurons (Fig. 1B-i). Bupivacaine caused morphological damage to the neurons and reduced axon growth, followed by cell death (Fig. 1B-iis). The high toxicity of bupivacaine towards PC12 cells, compared to 4T1 cells, allows specificity towards neurons.

### Bupivacaine lipid nanoparticles

Encapsulating bupivacaine in nanoparticles reduces systemic side effects and improves tumor targeting (*11, 17*). Bupivacaine was remotely loaded, using a transmembrane ammonium sulfate gradient (*17*), into 100±20 nm liposomes composed of hydrogenated soybean phosphatidylcholine (HSPC), cholesterol and 1,2-distearoyl-sn-glycero-3-phosphoethanolamine-N-methoxy-polyethylene glycol 2000 (DSPE-PEG2000), in molar ratios of 55:40:5, respectively (Fig. 1C-E). The encapsulation efficiency reached ∼60% (Fig. S2), being stable at 4 °C for 12 days, with a maximal release of 4% ± 1 (Fig. S1).

### Nanoparticle’s lipid composition affects neuronal uptake

To study the effect the nanoparticle’ lipid composition has on neuronal uptake we compared six different compositions of phosphatidylcholine (PC)-liposomes, varying in their lipid tail length and PEG moiety (Fig. 1F). PC and cholesterol-based liposomes were chosen as candidates for neuronal delivery, as these are the main lipid components in the membranes of neurons (*19*). Phospholipids with shorter fatty acid chains (and a resultant lower phase transition temperature, Tm) achieved greater uptake by PC12 cells. Specifically, the uptake of 2-dimyristoyl-sn-glycero-3-phosphocholine (DMPC, 14:0C lipid chain) liposomes was ∼2-fold higher than 1,2-dipalmitoyl-sn-glycero-3-phosphocholine (DPPC, 16:0C), and ∼3-fold higher than HSPC (18:0C) (*p*<0.0001). Interestingly, PEG, a common additive to nanoparticles used for stabilizing formulations and increasing circulation time (*20*), improved neuronal uptake of HSPC and DPPC liposomes, yielding ∼2-fold higher than liposomes without PEG (*p*<0.0001). Previous studies reported that PEG can improves tissue penetration (*21-23*), however, at the cell-level these observations are dependent on the cell-type. DMPC and DPPC liposomes displayed higher bupivacaine release rates at 37 °C compared to HSPC liposomes (*p*<0.05, from the third day) (Fig. 1G). Therefore, despite their superior cellular uptake, HSPC was used as the main membrane composition of liposomes in order to assure a stable formulation with a greater drug retention at physiological conditions (37 °C) (*24*).

### Cancer cells stimulate neurite growth through cytokine secretion

To examine the effect different cancer cell-lines have on nerve cells, PC12 cells were co-cultured with either 4T1 or pancreatic ductal adenocarcinoma (PDAC, KPC cell-line). Cancer cells stimulated neuronal PC12 differentiation and neurite growth, compared to the untreated control (*p*<0.0001), having a similar neurite stimulating effect as in neurons treated with nerve growth factor (NGF, Fig. 2A,B). Morphometric analysis showed that 37.1% ± 16.7 and 42.6% ± 12.9 of the cells were differentiated when PC12 cells were co-cultured with 4T1 or KPC cells, respectively. The average neurite number per cell was 1.8 ± 0.4 or 3.5 ± 1.0, respectively, with 4T1 or KPC (Fig. 2B). Thereby, neuronal differentiation was stimulated by the presence of both cancer cell-lines, 4T1 and KPC cells. The promotion of neurite growth by cancer cells was attributed to the secretion of neurotrophic factors (*25, 26*). Here we found that 4T1 cells secret chemokines C-C motif ligand 5 (CCL5), granulocyte colony-stimulating factor (GCSF), and chemokine C-C motif ligand 2 (CCL2) (Fig. 2C), that increase neurite growth and neural invasion (*25, 27-29*). Cancer cells promoted neurogenesis (*2*) in a multifactorial signaling system, which included the secretion of neurotrophic factors (*25, 26*) and cytokines.

**Figure 2.**
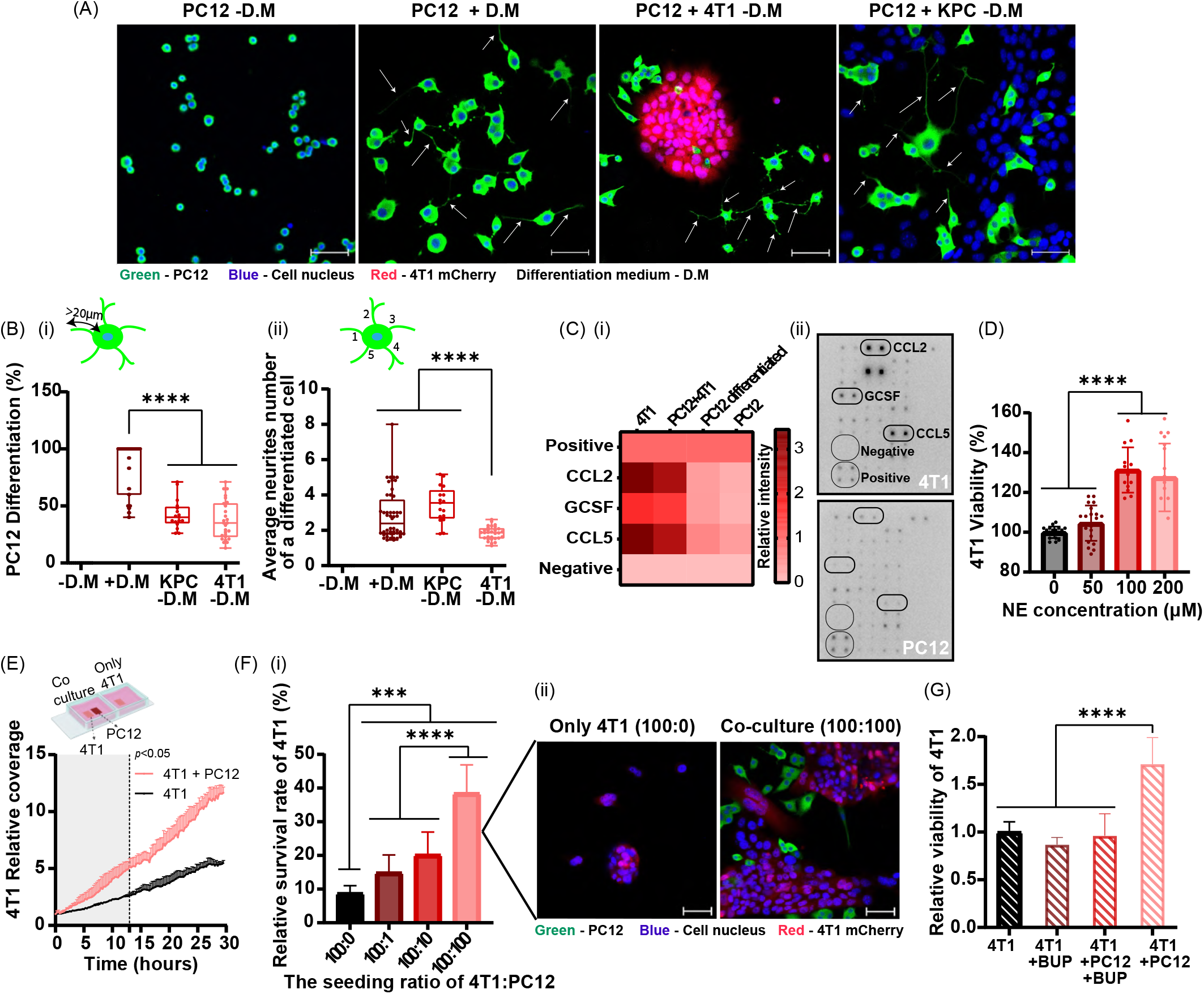
Nerve cancer cross-talk promotes neurite growth and cancer cells proliferation, migration and survival. *Cancer cells stimulate neurite growth through cytokine secretion (****A-C***). Confocal imaging of PC12 cells co-cultured with breast cancer 4T1 or pancreatic ductal adenocarcinoma (KPC) cells over 72 hours testing differentiation and neurite growth. Cell differentiation media (D.M) was supplemented with nerve growth factor only in the control group. Cell’ nuclei were labeled with Hoechst (blue). PC12 neurites are highlighted with arrows (**A**, scale bar is 50µm). Morphometric analysis of the co-culture indicates the percentage of differentiated cells (**B-i**), and the average number of neurites extending from the cell soma (**B-ii**). Cancer cells secrete cytokines that promote neurites outgrowth, identified by a cytokine antibody array (**C-i**) and presented as a heatmap (**C-ii**). *Neuronal cells enhance cancer cell proliferation, migration and survival (****D-G***). Effect of norepinephrine (NE) on 4T1 proliferation after 24 hours, normalized to the untreated group (**D**). The effect of PC12 on 4T1 migration; measured by time-laps imaging and analyzed using IMARIS, normalized to the beginning coverage (**E**). 4T1 cells survival under starvation conditions after 96 hours, with and without PC12 cells in the culture, measured using an InCell analyzer (**F-i**) and confocal microscopy (**F-ii** scale bar is 50µm). 4T1 cells were seeded with or without PC12 cells for 72 hours in starvation media, and then 0.5mg/ml bupivacaine (BUP) was added for 6 hours. Viability values are normalized to the cell count of 4T1 alone (**G**). Results of (B, D, F) (three independent repetitions preformed in at least 5 replicates) and (E,G) (two independent repetitions preformed in at least 2 replicates) are presented as mean±SD. One-way ANOVA with adjusted *p* value in multiple comparisons tests was used for statistical analysis of (B, D-F); \*\*\**p*<0.001, \*\*\*\**p*<0.0001

### Neuronal signaling promotes cancer cell proliferation, migration and survival

Nerve signaling is conducted through neurotransmitter secretion (*3*), such as norepinephrine (NE). Increasing concentrations of NE were used to evaluate its influence on cancer cells. 4T1 cells proliferation increased by ∼30% (*p*<0.0001) when treated for 24 hours with 100 µM and 200 µM of NE, compared to the untreated group (Fig. 2D). The effect of the presence of neurons on cancer cell migration and proliferation was monitored. For this, triple-negative 4T1 breast-cancer cells were seeded alone or together with neuronal PC12. Once 4T1 cells were co-cultured with PC12 cells, the migration and proliferation of 4T1 cells increased compared to 4T1 alone. (Fig. 2E). In addition, the co-culture of 4T1 with PC12 cells contributed to cancer cell survival under starvation conditions, in a concentration dependent manner (Fig. 2F). When 4T1 were co-cultured with PC12 cell at a 1:1 or 10:1 ratio, the survival rate increased by 4.4-fold and 2.3-fold, respectively, compared to the survival of 4T1 that were seeded alone (*p*<0.0001). Accordingly, nerve cells promote breast cancer cell proliferation, spread and survival (*1, 26*).

### Targeting neurons with bupivacaine reduces cancer cell viability

To suppress nerve/cancer interactions bupivacaine was used as a neurotoxic agent. In PC12/4T1 co-culture, 4T1 cell viability was ∼2-fold higher compared to 4T1 cells seeded alone (*p*<0.0001) (Fig. 2G). However, addition of bupivacaine in neurotoxic concentration to the co-culture medium annulled this effect, and cancer-cell viability was decreased ∼2-fold (*p*<0.0001), similarly to 4T1 seeded alone treated with bupivacaine or untreated 4T1. Bupivacaine caused neuron-specific toxicity, which inhibited nerve/cancer cell interaction, followed by a decrease in cancer cell growth. These *in-vitro* findings underscore the role of nerve cells in cancer development, and the potential of targeting the neurons within the tumor tissue with bupivacaine in order to inhibit cancer progression.

### The uptake of 100-nm liposomes by neurons and cancer cells

We compared the liposomal uptake in PC12 and 4T1 cells (Fig. 3). Flow cytometry analysis demonstrated an increased liposomal uptake over time by 4T1 and PC12 cells (Fig. 3B). After 1 and 12 hours, liposomal uptake by 4T1 cells increased from 9.5%±4.3 to 94.9%±3.2, respectively, while the liposomal uptake by PC12 cells increased from 11.2%±5.0 to 72.6%±12.9. Confocal microscopy analysis demonstrated that liposomes accumulated after 1 hour in the neuronal bodies, and after 4 hours they were detected also along the axons (Fig. 3A). The liposomes appeared to be fused with the PC12 cell membrane, while in 4T1 cells liposomes were clustered in the cytoplasm (*30, 31*). We further examined the liposomal uptake in cortical primary neurons cells, where liposomes accumulated in both neuronal cell body and fibers after 2 hours (Fig. 3C). The liposomes were also taken up by primary astrocytes and microglia, but to a lower extent than the primary neurons (Fig. 3D). Our findings emphasize the potential of nanotechnology in neurobiology research, such as neuronal targeting or recovery.

**Figure 3.**
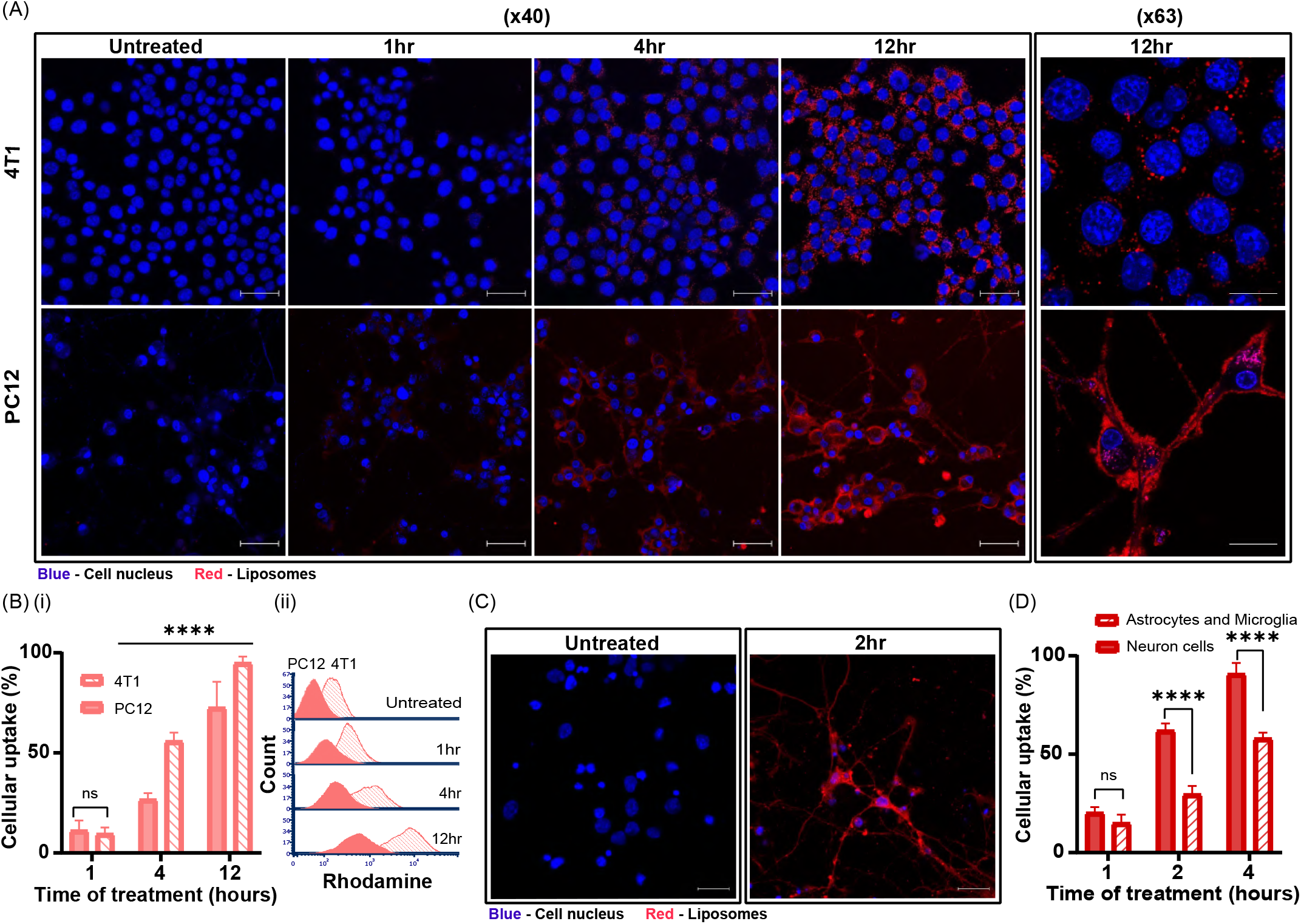
The uptake of 100-nm liposomes by neurons cancer cells. The cellular uptake of 100 nm Rhodamine-labeled liposomes by PC12 and 4T1 cells over time, was observed using confocal microscopy (**A**, scale bar is 50µm for X40 magnification and 20µm for X63 magnification) and flow cytometry analysis (**B-i**) and histogram (**B-ii**). Cellular uptake of liposomes by primary cortical neurons and astrocytes and microglia, were imaged using confocal microscopy (**C**, scale bar is 20µm) and compared flow cytometry (**D**) analysis. Results of B (three independent repetitions preformed in at least 3 replicates) and D (two independent repetitions preformed in at least 3 replicates) are presented as mean±SD. One-way ANOVA with adjusted *p* value in multiple comparisons tests was used for statistical analysis; \*\*\*\**p*<0.0001.

### Neurons are integral in breast cancer tumors

Identifying neurons in 4T1 breast cancer *in vivo* model was conducted using two different immunohistochemistry specific neuronal markers, anti-β III Tubulin and anti-PGP9.5 (Fig. 4A, B). The nerve trunk, as well as neuronal cell bodies and axons, were detected and distributed throughout the non-necrotic areas of the tumor tissue. Quantitative flow cytometry analysis of tumors dissociated into single cell suspensions after staining for adrenergic neurons (using anti-tyrosine hydroxylase), demonstrated that 3.6% ± 0.6 of the total tumor cell population were neurons (Fig. 4C). Nerves inside 4T1 tumors tissue demonstrates their active innervation into the tumor tissue and their potential involvement in tumor development (*2, 26*).

**Figure 4.**
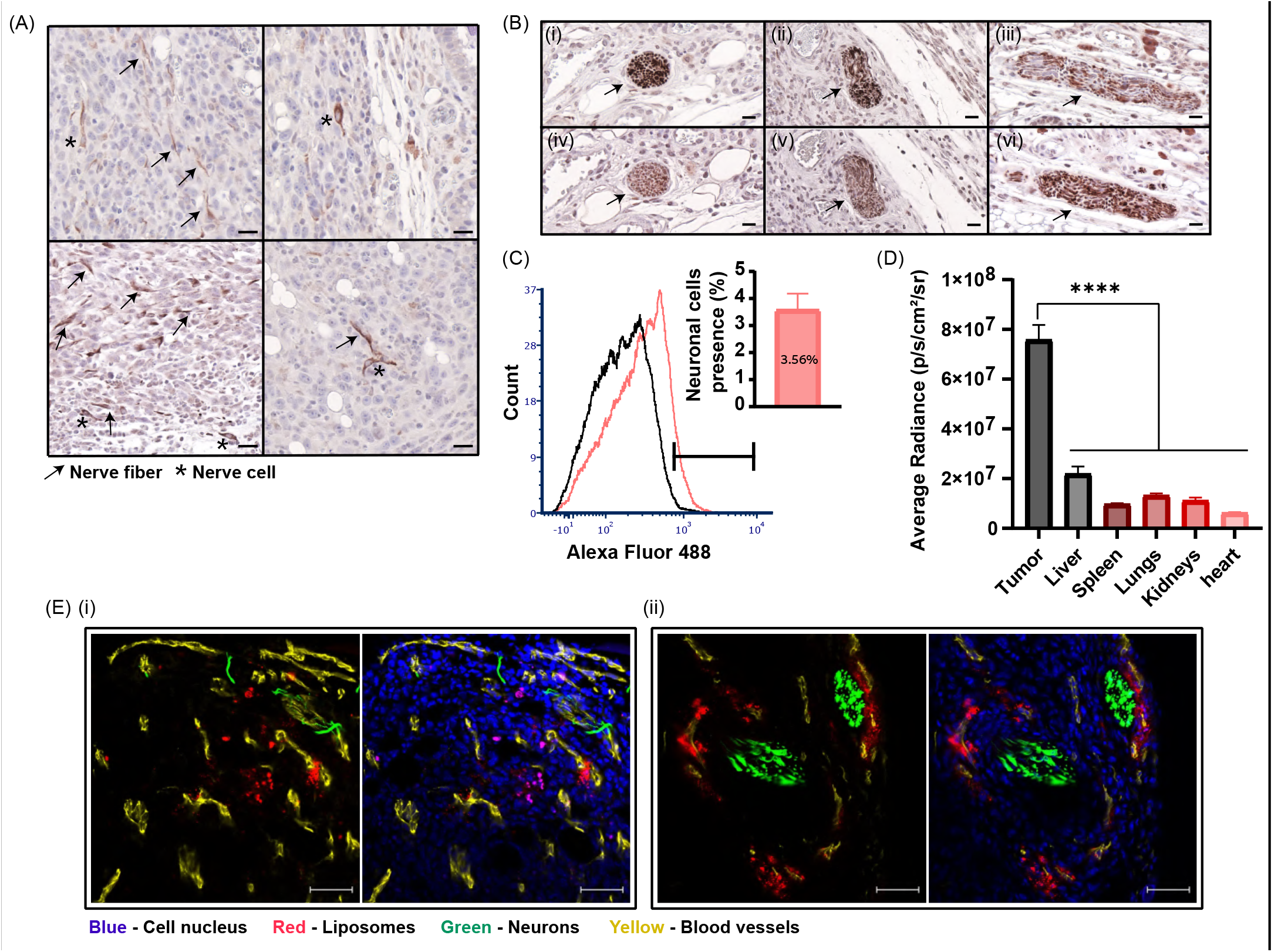
Biodistribution of 100-nm liposomes to orthotopic breast cancer tumors and their delivery to tumor neurons. Detection of nerve fibers and cells in 4T1 tumor tissue sections was performed using immunohistochemistry staining of anti-beta III Tubulin. Nerve fibers and cells are represented by arrows and stars, respectively (**A**, scale bar is 20µm). Co-immunostaining of serial sections (i, iv and ii, v and iii, vi) of the 4T1 tumor tissue with anti-PGP9.5 and anti-beta III Tubulin demonstrated nerve trunk presence in breast cancer (**B**, scale bar is 20µm). Detection of adrenergic neurons with anti-Tyrosine Hydroxylase in 4T1 tumor tissue was performed using flow cytometry (**C**). Rhodamine-labeled liposomes were injected intravenously to mice bearing orthotopic 4T1 tumors. 24-hours post administration nanoparticle’ biodistribution to different tissues was quantified using IVIS analysis (**D**). Liposomal accumulation within the tumor tissue was also demonstrated by fluorescent histology (**E**, scale bar is 50µm). Nerve infiltration (beta III Tubulin+) and liposome accumulation are associated with blood vessels presence (CD31+). Results of (C) (n=10) and (D) (n=3) are presented as mean±SEM. One-way ANOVA with adjusted *p* value in multiple comparisons tests was used for statistical analysis; \*\*\*\**p*<0.0001.

### Nanoparticle delivery to tumor neurons

We explored the capacity of liposomes to target neurons within the tumor tissue. Rhodamine-labeled liposomes were injected intravenously to mice bearing orthotopic 4T1 tumors. 24-hours post administration, the tumors were excised and the liposomal biodistribution was assessed using *ex-vivo* IVIS imaging (Fig. 4D) and fluorescent histology (Fig. 4F). Increased liposomal accumulation at the tumor tissue was recorded compared to healthy tissues (Fig. 4D). Histological examination of the tumor tissue demonstrated nerve infiltration into the tumor and liposome distribution with the tumor neurons (Fig. 4F, S4B). These findings are attributed to the infiltration of sympathetic nerve fibers into the tumor with its vasculature (*2, 7*), and to the preferential delivery of nanomedicines to the tumor microenvironment through its blood supply (*11*). Therefore, the intravenously treatment of bupivacaine loaded liposomes enables the specific targeting of neurons within the tumor tissue.

### Nanoparticle bupivacaine inhibits tumor growth and metastases

Liposomal bupivacaine (L-BUP) was administered to mice bearing orthotopic 4T1 tumors (Fig. 5A). The median lethal dose (LD_50_) of bupivacaine in mice is between 6 mg/kg to 8 mg/kg per body weight (*32*). Therefore, when injecting a high concentration of bupivacaine, it was of paramount importance to use liposomes rather than that free form of the drug. L-BUP was administered at two different concentrations, 10 mg/ kg-body-weight and 3.5 mg/kg. To compare the efficacy of L-BUP with other common treatment, liposomal doxorubicin (L-DOX) was administered at 4 mg/kg. No signs of toxicity were observed during the treatment with L-BUP and mice’s weight increased gradually similarly to the untreated group (Fig. S5). The higher dose of 10 mg/kg L-BUP inhibited tumor growth similar to L-DOX treatments (Fig. 5B, C), while L-BUP at 3.5 mg/kg did not affect tumor progression (*p*<0.05). Therefore, a higher dose of bupivacaine is needed to achieve a therapeutic effect.

**Figure 5.**
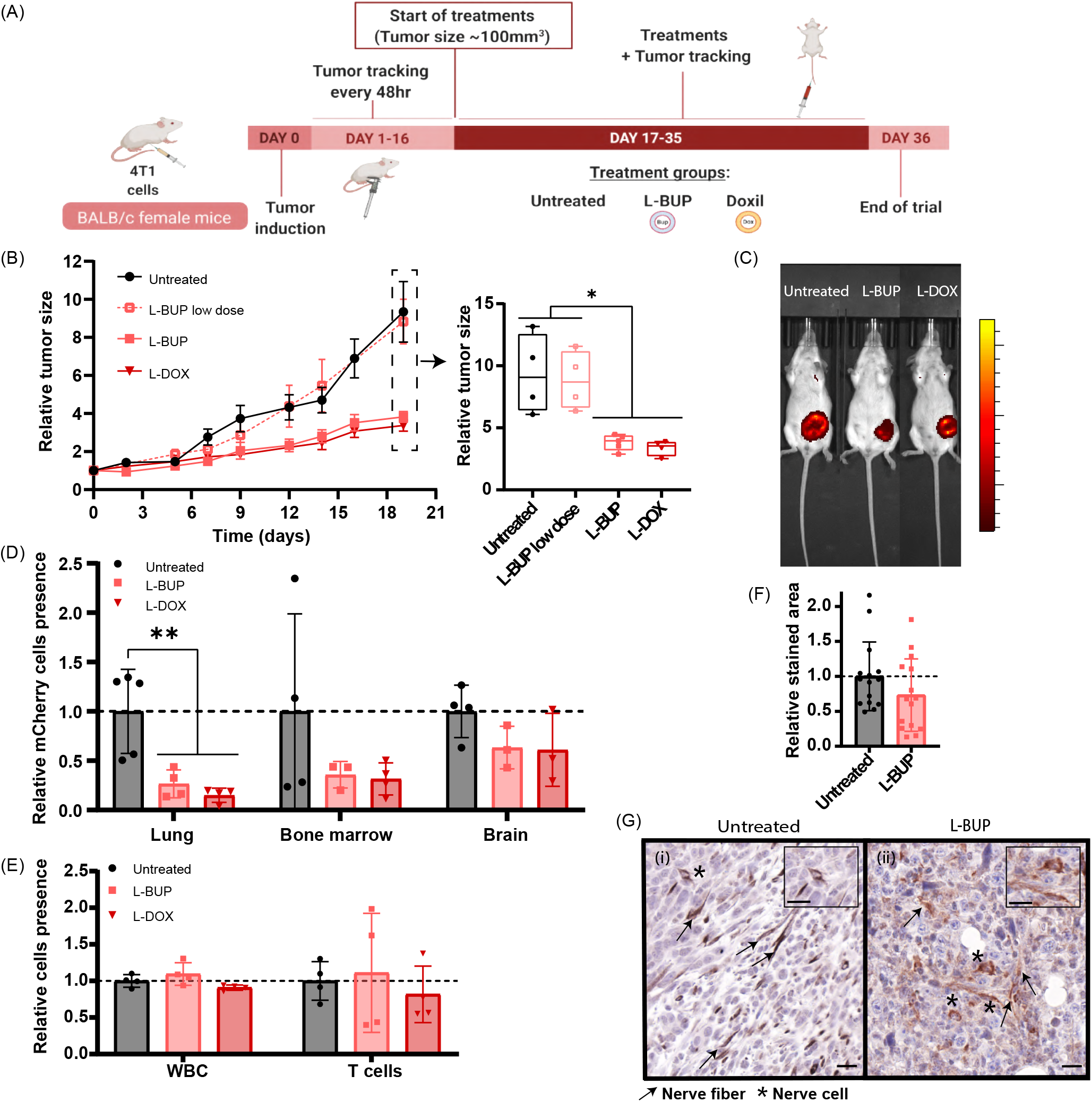
Liposomal bupivacaine inhibits tumor growth and metastases. Once the tumors reached 100mm^3 liposomal bupivacaine (L-BUP) at a high dose (10mg/kg) or low dose (3.5 mg/kg), or liposomal doxorubicin (L-DOX) (4 mg/kg-body weight, positive control), were administered intravenously. (**A**). Tumors were sized every two/three days (**B**, normalized to day 0) and were imaged by IVIS at the end of the trial (**C**). L-BUP treatment reduced metastases formation in lung (p<0.01), bone marrow (p<0.6), and brain (p<0.8), compared to the untreated group. Metastases was detected by dissociating the tissue into a single-cell suspension and then detecting 4T1-mCherry using flow cytometry (**D**, normalized to the untreated group). Moreover, the flow cytometry analysis indicated there was no change in the level of total white blood cell (CD31+) and T-cell (CD45+/CD3+) count in the tumor tissue in the different treatments (**E**, normalized to the untreated group). Nerve density in 4T1 tumor tissue sections, with and without L-BUP treatment, was evaluated by immunohistochemistry staining and analysis of anti-beta III Tubulin (**F**, normalized to the untreated group). Nerves structure was damaged due to L-BUP treatment (G-ii) compared to the untreated group (G-i). Nerve fibers are represented by arrows, and nerve cells are represented by stars (**G**, scale bar is 20µm). Results of (B) (4≤n≤6) are presented as mean±SEM. Results of (D-F) (3≤n≤5) are presented as mean±SD. Two-tailed unpaired Student’s t-test was used for statistical analysis of (B and F) and One-way ANOVA for (D, E) with adjusted *p* value in multiple comparisons tests; \**p*<0.05 and \*\**p*<0.01.

Five weeks post tumor induction mice were sacrificed, and the lungs, brain, and bone marrow were resected and dissociated into single cells for quantitative detection of metastasis by flow cytometry (Fig. 5D). L-BUP inhibited metastases in the lung (*p*<0.01), bone marrow (*p*<0.6), and the brain (*p*<0.8) as affectively as L-DOX and in comparison to the untreated group.

The effect of the different treatments on the infiltration of immune cells (CD45+) and T cells (CD45+/CD3+) was examined using flow cytometry (Fig. 5E). All the treatment groups had a similar immune cell profile, suggesting that the observed therapeutic effect of L-BUP is related to non-immune mechanisms, such as the involvement of the nervous system, and not as a result of immune response activation.

The difference in nerve density within the tumor tissue between L-BUP and the untreated group was examined by immunohistochemistry staining of anti-β III Tubulin in different sections. Reduced tumor’ nerve density was recorded in the L-BUP treatment group (*p*≈0.2) (Fig. 5F) and the nerves were morphologically damaged (Fig. 5G), similarly to neurons exposed to the bupivacaine *in vitro* (Fig. 1B-ii).

Overall, we used bupivacaine-loaded liposomes in order to suppress nerves within the tumor microenvironment and demonstrated that L-BUP inhibited tumor growth and metastatic dissemination without causing any toxic effect nor sever immunogenic response.

## Discussion

Nerves actively infiltrate the tumor microenvironment, and promote tumor development and progression through neurotransmitter secretion. Their recruitment is driven by the cancer cells, through the secretion of neurotrophic factors and cytokines (*1, 25*). Here we present the use of nanotechnology for cancer-neural therapy. Interruption of nerve to cancer cross-talk using liposomal bupivacaine was achieved and consequently inhibited cancer development. In co-culture of PC12 and 4T1, free bupivacaine reduced the viability of the neuronal cells. We developed a liposomal bupivacaine formulation to reduce its systemic adverse effects and improve tumor uptake. 100-nm liposomes administered intravenously to mice bearing orthotopic 4T1 tumors were distributed throughout the tumor and specifically with the tumor neurons, curbing tumor growth. Hence, L-BUP may be leveraged as an alternative tool for treating breast cancer. In addition, non-opioid liposomal bupivacaine (*32*) may be a new strategy for dealing with cancer pain owed to sensory nerves in the tumor microenvironment (*1*).

Overall, we demonstrate the collaborative interactions between nerves and cancer, and the potential of analgesic nanotechnology to supress these interactions. This study suggests that targeting nerves in the tumor tissue using non-opioid anesthetic nanoparticles is a potential new clinical approach that can improve breast cancer therapy.

## Acknowledgments

This project received funding from the European Union’s Horizon 2020 research and innovation program under the grant agreement No 680242-ERC-[Next-Generation Personalized Diagnostic Nanotechnologies for Predicting Response to Cancer Medicine]. The authors also acknowledge the support of Israel Innovation Authority for the Nofar Grant (67967), the Israel Science Foundation (1778/13, 1421/17); The Israel Ministry of Economy for a Kamin Grant (52752, 69230); the Israel Ministry of Science Technology and Space – Office of the Chief Scientist (3-11878); the Israel Ministry of Science, Technology & Space (3-16963; 3-17418); the Israel Cancer Association (2015-0116); Leventhal 2020 COVID19 Research Fund (ATS #11947), the German-Israeli Foundation for Scientific Research and Development for a GIF Young grant (I-2328-1139.10/2012); the European Union FP-7 IRG Program for a Career Integration Grant (908049); the Phospholipid Research Center Grant (ASC-2018-062/1-1); the Louis family Cancer Research Fund, a Mallat Family Foundation Grant; The Unger Family Fund; a Carrie Rosenblatt Cancer Research Fund, the Technion Integrated Cancer Center (TICC), the Russell Berrie Nanotechnology Institute, the Lorry I. Lokey Interdisciplinary Center for Life Sciences & Engineering. A. Schroeder acknowledges the Alon and Taub Fellowships. Maya Kaduri and Mor Sela wish to thank TEVA Pharmaceuticals - NFBI - The National Forum for Bio-Innovators for the Doctoral Fellowship. Maya Kaduri wishes also to thank to the Technion Integrated Cancer Center (TICC) Rubinstein scholarship, and Robert B. Kalmansohn Fellowship Fund in the Emerson Life Sciences Building. Maria Poley wishes to thank the Israeli Ministry of Science and Technology for the Shulamit Aloni Doctoral Fellowship. Hanan Abumanhal wishes to thank the Baroness Ariane de Rothschild Women Doctoral Program. Dr. Alberto Ouro wishes to thank the Basque Country Government for the Post-Doctoral Fellowship. The authors also wish to thank professor Asya Rolls from the Faculty of Medicine, Technion, and Tamar Ben-Shaanan from University of California, San Francisco, USA, for their professional input and helpful discussions that greatly contributed to the paper. Images in this paper were created with BioRender.com and Adobe Illustrator. The assistance of Mrs. Irina Davidovich from the Technion Center for Electron Microscopy of Soft Matter for the Cryo-TEM measurements was greatly appreciated.

## Materials and Methods

### Bupivacaine loaded liposomes preparation

Lipid mixture of HSPC or DPPC or DMPC (Lipoid, Ludwigshafen, Germany), cholesterol (Sigma-Aldrich, Rehovot, Israel) and DSPE-PEG2000 (Lipoid, Ludwigshafen, Germany), in molar percentages of 55:40:5 were dissolved in warm absolute ethanol. Bupivacaine was actively loaded into the liposomes using the ammonium sulfate gradient method(*17*). Shortly, once all the lipids were completely dissolved in ethanol, the lipid suspension was added into 250 mM ammonium sulfate solution to reach a final concentration of 50 mM total lipids, which corresponds to 1.5×10^13^ liposomes/ml. To obtain homogenous 100 nm liposomes, the liposome mixture was extruded through 400, 200, 100 and 80 nm pore-size polycarbonate membranes (Whatman, Newton, MA, USA) using a Lipex extruder (Northern Lipids, Vancouver, Canada) at a temperature that is above the Tm (70°C or 60°C or 50°C for HSPC, DPPC and DMPC respectively). The liposomes solution was dialyzed against saline (150 mM NaCl at pH 5.5) (1:1000 volume ratio) using a 12-14 kD dialysis membrane (Spectrum Laboratories, Inc., USA) at 4°C and exchanged three times (after 1, 4 hours and overnight). For active loading, bupivacaine hydrochloride (Sigma-Aldrich, Rehovot, Israel) was dissolved in saline at pH 5.5 and added to the ammonium sulfate liposomes to reach a final concentration of 5 mg/ml. The mixture was incubated at 800 rpm for 1 hour at a temperature that is above the Tm. The non-encapsulated bupivacaine was removed by dialysis against saline in the same manner as described above.

Liposomes size analysis, which includes mean diameter (nm) and particle size distribution (PDI) measurements, were carried out by dynamic light scattering (DLS) using a Zetasizer Nano ZSP (Malvern, UK). Particles concentration (liposomes/ml) measurements were carried out using a Zetasizer Ultra (Malvern, UK).

### Rhodamine-labeled liposomes preparation

Rhodamine (Avanti Polar Lipids, Alabaster, AL, USA) labeled liposomes were also prepared at the same method of ethanol injection as described above. Shortly, 16:0 Liss Rhod PE was added to the lipid mixture at 0.1% molar ratio. As for the buffers, the lipid mixture was injected into a Phosphate Buffer Saline (PBS; Sigma-Aldrich, St. Louis, USA). To compare between different liposomal compositions, liposomes composed of DMPC (Lipoid, Ludwigshafen, Germany) and DPPC (Lipoid, Ludwigshafen, Germany) were also prepared in the same molar ratios. As for the liposomes without DSPE-PEG2000, the composition was HSPC:cholesterol in molar percentages of 60:40. In addition, when preparing the DMPC-liposomes, 14:0 Liss Rhod PE was used instead of 16:0.

### Doxorubicin loaded liposomes preparation

Doxorubicin (DOX, TEVA Israel) was actively loaded into 100 nm liposomes using the ammonium sulfate gradient method (*33*). The dissolved lipid mixture of 55:40:5 of HSPC, cholesterol and DSPE-PEG2000 was injected into 120 mM ammonium sulfate solution to reach a final concentration of 50 mM total lipids, which corresponds to 1.5×10^13^ liposomes/ml. The liposome mixture was extruded as described above, and then dialyzed against 10% w/w sucrose (1:1000 volume ratio) with three exchanges during 24 hours. For active loading, DOX was dissolved in 10% w/w sucrose and added to the ammonium sulfate liposomes to reach a final concentration of 2 mg/ml. The mixture was incubated in 65°C at 800 rpm for 1 hour. The non-encapsulated DOX was removed by dialysis against 10% w/w sucrose in the same manner as previously.

### Cryogenic transmission electron microscopy

Cryogenic transmission electron microscopy (cryo-TEM) imaging was performed using a FEI (Thermo Fisher Scientific) Talos 200C high-resolution TEM (Technion Center for Electron Microscopy of Soft Matter, at the Wolfson Department of Chemical Engineering). Specimen preparation was carried out at controlled conditions of 25°C and 100% relative humidity in a Controlled Environment Vitrification System (CEVS). A drop of diluted liposomal bupivacaine (0.5mM) was placed on a carbon-coated perforated polymer film, supported on a 200 mesh TEM grid, mounted on a tweezer. The drop was thinned into a film with a filter paper-covered metal strip, and then was quickly plunged into liquid ethane at its freezing point (−183°C). The grid was transferred under controlled conditions into Gatan 626 (Gatan, Pleasanton, CA) cryo-holder, and imaged at -175°C. Images were recorded digitally by an FEI Falcon III, highly sensitive direct-imaging camera. A Volta “phase-plate” was used to enhance image contrast.

### Lipids and bupivacaine concentration measurements

High Performance Liquid chromatography (HPLC, 1260 infinity, Agilent Technologies, Santa Clara, CA) equipped with an ELSD and UV detectors was employed to quantify the lipid composition of the liposomes and the loaded bupivacaine concentration. Absorption was measured at a wavelength of 230 nm. The separation was achieved using Luna C18 column, 5 mm, 100 Å (Phenomenex LTP, Aschaffenburg, Germany). The mobile phase consisted of three solutions; A 0.2% Trifluoroacetic acid (TFA) in water, B 0.2% TFA in methanol, and 0.2% TFA in isopropanol. Bupivacaine separation was achieved by starting conditions of 80% A and 20% B, followed by a linear gradient up to 10% A and 90% B for 15 min, at 40°C with constant flow rate of 1.5 ml/min. Then, lipids separation was completed by keep using the last solvent composition for 5 more minutes, and then by gradually changing the composition to 80% B and 20% C within 22 minutes at 40°C with constant flow rate of 2.2 ml/min. The solvent composition gradually returned to the opening conditions within 18 minutes. ELSD settings were adjusted to 1.6 SLM of the inert gas flow, 60°C as the nebulizer temperature and 80°C as the evaporator temperature. Sample was prepared by dilution the liposomes 1:20 in methanol and injection volume was 10 μl. Calibration curves were obtained according to the peak areas in the chromatograms and used for quantification of analyzed samples concentration.

To verify the loaded bupivacaine concentration in another method, the liposomes were diluted 50-fold with ethanol (to release the encapsulated bupivacaine) and measured at a wavelength of 230 nm by plate reader (Tecan, Mannedorf, Switzerland). Calibration curve was prepared by diluting bupivacaine hydrochloride together with the ammonium sulfate liposomes in ethanol.

The encapsulation efficiency (60%) was determined as the percentage of the final loaded bupivacaine concentration according to HPLC (3 mg/ml), to the initial bupivacaine concentration that was used for the remote loading (5 mg/ml). The drug-to-lipid mole ratio in the nanoparticles (0.3) was calculated by dividing the loaded bupivacaine concentration (9.2 mM) by the final lipids concentration (30.5 mM), as was determined by HPLC (Fig. S2).

### Bupivacaine release profile

Liposomal bupivacaine were dialyzed against saline (1:10 volume ratio) at 4°C or 37°C at 200 rpm. The level of free bupivacaine in the extra-liposomal buffer was quantified at the desired time intervals by measuring 100 µl of sample at plate reader in 230 nm wavelength. The sample was then returned back to the incubation until the next measurement. The percentage of bupivacaine released was determined by the ratio of free bupivacaine released to total encapsulated bupivacaine concentration Cell culture

### Cell culture

All cells were cultured in 37°C in a humidified atmosphere and 5% CO_2_ in air, and every 2 days a fresh medium was added to the cells.

Triple negative breast cancer cell line (4T1) (ATCC) were grown in Roswell Park Memorial Institute medium 1640 (RPMI 1640, Sigma-Aldrich, Rehovot, Israel) supplemented with 10% v/v of fetal bovine serum (FBS, Biological Industries, Beit Haemek, Israel), 1% v/v Penicillin-Streptomycin solution (100 1U/ml of Penicillin G Sodium Salt and 100 µg/ml of Streptomycin Sulfate) (Pen-Strep), and 1% v/v L-Glutamine (Biological Industries, Beit Haemek, Israel). 4T1 cells expressing mCherry were developed (*34*) and kindly provided by the Ronit Satchi-Fainaro (Cancer Research and Nanomedicine Laboratory, Tel Aviv University). Puromycin was added to cell medium in a final concentration of 10 µg/ml.

KPC cells were established in the laboratory of Surinder K. Batra. KPC cells were LSL KrasG12D/+;LSLTrp53R172H/+ of pancreatic carcinomas, along with inactivation of the ;Pdx-1-Cre (KPC) transgenic mice (*35*). Cells were cultured in Dulbecco’s modified Eagle’s medium (DMEM; Sigma-Aldrich) supplemented with 10% v/v of FBS, 1% v/v Pen-Strep and 1% v/v L-Glutamine.

PC12 cells (ATCC) were kindly provided by Orit Shefi (Neuro-engineering and Regeneration laboratory, Faculty of Engineering, Bar-Ilan). Cells were cultured in suspension in the RPMI medium supplemented with 10% v/v horse serum (HS, Biological Industries, Beit Haemek, Israel), 5% v/v FBS, 1% v/v Pen-Strep, 1% v/v L-glutamine, and 0.2% v/v amphotericin B (Biological Industries, Beit Haemek, Israel). To induce their differentiation, PC12 cells were seeded on collagen type l (Corning Inc., NY, USA.) coated plates in serum reduced media (1% HS) supplemented with 50 ng/mL of murine β-NGF (Peprotech, Israel).

Primary cultures of neonate rat pups (P0–P4) cortical neurons were established and kindly provided by Shai Berlin (Department of Neuroscience, The Ruth and Bruce Rappaport faculty of Medicine, Technion-Israel Institute of Technology), and were produced as was described previously (*36*). These cells were used for confocal analysis of liposomal uptake. Primary cultures of E13.5 or E16.5 mouse embryo cortical neurons were kindly provided by Dr. Yossef Buganim (Department of Developmental Biology and Cancer Research, Institute for Medical Research Israel-Canada, The Hebrew University-Hadassah Medical School), and were produced as was previously described(*37*). These cells were used for flow cytometry analysis of liposomal uptake. Cells were plated into 6-wells plate coated with poly-l-lysine (0.1 mg/ml). Neurons were grown in neurobasal medium (Gibco, Ireland) plus B27 supplement (Gibco, Ireland) and GlutaMAX (Gibco, Ireland) until day 5, then medium was replaced with Neurobasal medium plus B27 with Cytosine β-D-arabinofuranoside (Ara-C) at 10 µM, to avoid the proliferation of glial cells, and without GlutaMAX for another 11 days, up to 14 days *in vitro* (DIV), where neuronal maturity is considered to be reached.

### Breast cancer model

Eight to ten week-old BALB/c female mice (Harlan laboratories, Jerusalem, Israel) were used as breast cancer animal models. 50 µl of 4×10^5^ 4T1 cells were injected subcutaneously into the fourth mammary fat pad to obtain primary tumor model. All animal experiments were approved by, and in compliance with, the Inspection Committee on the Constitution of the Animal Experimentation at the Technion, Israel Institute of Technology.

### Confocal imaging

Confocal microscopy (LSM-700, Zeiss, Germany) was employed to examine several experiments such as co-culture studies, liposomal uptake etc. For tissue culture imaging, PC12 cells were stained over-night with rabbit polyclonal anti-Tyrosine hydroxylase (Abcam, Cambridge, UK), following a staining with donkey polyclonal anti-rabbit IgG H&L conjugated Alexa Fluor 488 (Abcam, Cambridge, UK). Antibodies were diluted by 1:1000 in a blocking serum. All cells were stained with 1 µg/ml Hoechst for nuclei labeling. Acquisition was performed using the ZEN software and applying the 405, 488, 555 and 639 nm lasers.

### Imaging and morphometric analysis of PC12

In order to study the effect of cancer cells on nerve cells, PC12 cells were co-cultured with 4T1 mCherry and KPC. Cells were seeded in an optical µ-slide 8-well Collagen IV coated plate (ibidi, Madison, WI) in serum reduced media for 72 hours, at a density of 3×10^4^ cells/well for PC12 cells and at a density of 1.5×10^4^ cells/well for 4T1 and KPC. As a positive control to neuronal differentiation, 24 hours from seeding NGF was added to the medium of some of the PC12 cells that were seeded alone(*18*). After 72 hours from seeding, PC12 cells were stained and imaged by confocal microscope. All the confocal images were taken to quantitative morphometric analysis by the IMARIS software, which enables semi-automatic tracing of neurites and length measurements. The analysis included two accepted morphological parameters at the single-cell level: (i) the percentage of differentiated cells out of the total cell population, and (ii) the average number of neurites, which can be either axons or dendrites, extending from the cell body of a differentiated cell. A cell was regarded as differentiated when its neurite length exceeded 20μm from the nucleus. Morphological parameters and statistics were measured for at least 20 fields of 3 independent repetitions.

### Quantification of 4T1 cell confluence in PC12/4T1 co-culture

In order to study the effect of nerve cells on cancer cells, PC12 cells were co-cultured with 4T1 in serum reduced media for 96 hours. 4T1 mCherry cells were seeded in 24-well plates at a density of 7×10^4^ cells/well, and PC12 cells were seeded together with 4T1 in increasing density of 0, 7×10^2^, 7×10^3^ and 7×10^4^ cells/well. Each plate was scanned by GE InCell Analyzer2000 to obtain random images (13 fields per well, and 6 wells per each treatment) using DAPI and Texas Red channels. The plates were scanned 24 hours from seeding (was set as start-point) and again after 96 hours (was set as end-point). Before each scanning cells were stained with 1 µg/ml Hoechst for nuclei labeling. The images were analyzed using the InCell software to quantify the amount of nuclei of 4T1 mCherry cells. In order to determine the percentage of the survival rate of 4T1 mCherry cells, the obtained value for each well at the end-point of the experiment was divided by the value of the same well at the start-point.

A similar experiment was conducted with the presence of bupivacaine. PC12 cells were co-cultured with 4T1 mCherry in serum reduced media in 24-well plates at a density of 10×10^4^ cells/well for each type cell. After 72 hours from seeding, the medium was replaced to a fresh reduced medium without or with bupivacaine (0.5 mg/ml), and the cells were incubated for another 6 hours. Then cell nuclei were stained with 1 µg/ml Hoechst, and each plate was scanned by GE InCell Analyzer2000 using DAPI and Texas Red channels. The obtained images were analyzed using the InCell software to quantify the amount of 4T1 mCherry cells according to their intensity. To determine the relative viability of 4T1 mCherry cells, all of the values were normalized to the value of the untreated group (4T1 mCherry cells that were seeded alone).

### Cell viability assay

Cell viability was determined using a commercial 3-(4,5-dimethylthiazol-2-yl)-2,5-diphenyl tetrazolium bromide (MTT) viability assay (Sigma Aldrich, Rehovot, Israel). That assay was used to determine how highly 4T1 proliferation was stimulated by Norepinephrine (NE) bitartrate (Sigma Aldrich, Rehovot, Israel). 4T1 cells were seeded in 96-well plates at a density of 1×10^4^ cells/well. After 24 hours, the medium was replaced with serum reduced medium for another 24 hours. Then, the medium was replaced with fresh serum reduced medium supplemented with an increasing concentration of NE (0, 50, 100 and 200 µM), and the cells were incubated for 24 hours. Then, the medium was removed and the MTT assay was conducted.

MTT assay was also used to determine the toxicity of bupivacaine on 4T1 and PC12. 4T1 cells were seeded in 96-well plates at a density of 2.5×10^4^ cells/well two days before the experiment. PC12 cells were seeded separately in 96-well plates at a density of 3×10^4^ cells/well 6 days before the experiment, and treated two times with NGF to be fully differentiated. Both cell types were treated with an increasing concertation of bupivacaine (0, 0.5, 0.7 and 0.9 mg/ml) for 6 hours. Then, the medium was removed and the MTT assay was conducted. This experiment was also imaged by confocal microscopy, as described above.

### Migration assay

PC12 and 4T1 mCherry cells were seeded separately in 2 well silicone insert (ibidi, Madison, WI) on an optical µ-slide 8-well Collagen IV coated plate (ibidi, Madison, WI). As a control, 4T1 mCherry cells were seeded alone. PC12 cells were seeded at a density of 8.5×10^4^ cells/well in serum reduced media supplemented with NGF, and 24 hours later mCherry cells were seeded at a density of 3.5×10^4^ cells/well. 24 hours later, the media and the silicone insert were removed, the wells were filled with medium, and the plate was imaged by ZEISS Cell Observer SD for live imaging using brightfield and CY3 channel. The cells were monitored every 15 minutes, during 30 hours, and the images were analyzed by IMARIS software to obtain the area cover of 4T1 mCherry cells. All the values were normalized to the area cover measured at the beginning of the experiment.

### Cytokine kit array

PC12 cells were co-cultured with 4T1 for 72 hours. Cells were seeded in 6-well plate in serum reduced media at a density of 4×10^5^ cells/well for PC12 cells, and at a density of 2.5×10^5^ cells/well for 4T1. In addition, 4T1 cells were seeded alone at a density of 3×10^5^ cells/well. Cytokine profile of the conditioned medium was determined using Mouse Cytokine Antibody Array (Abcam, Cambridge, UK). The membranes were imaged by FUSION FX 6 (VILBER, France). Quantitative data was assessed using the ROI tool in Fiji software, and the obtained values were normalized to the positive control of each membrane.

### Evaluation of liposomal cellular uptake

PC12 cells were seeded 6 days before the experiment, and treated twice with NGF to be fully differentiated. PC12 and 4T1 were incubated for different time durations with Rhodamine-labeled liposomes in a final concentration of 0.5 mM total lipids, which corresponds to 1.5⨯10^11^ liposomes/ml. Then the culture media was removed and the cells were rinsed with PBS for three time, incubated with 0.25% trypsin-EDTA for 5 minutes, and centrifuged at 400Xg for 5 minutes. The cell pellet was resuspended in fresh medium. Cellular uptake was detected in the mCherry channel after acquisition of 10,000 single cells per sample using a flow-activated cell sorter (FACS; FACS Aria III, BD Biosciences). Analyses were conducted by FCS Express (De Novo software).

Primary neurons were incubated for different time durations with Rhodamine-labeled liposomes in a final concentration of 5 mM total lipids, which corresponds to 1.5×10^12^ liposomes/ml. After detaching and rinsing the cells as described above, cells were incubated in 10% FBS for 30 minutes at room temperature (RT). Subsequently, the cells were incubated with specific neuronal marker CD24-FITC Clone M1/69 antibody(*38*) (Stemcell Technologies, Canada) diluted 1:100 for 30 minutes on ice. Then the cells were rinsed with PBS for three time, and resuspended in PBS with 5% v/v FBS. Cellular uptake was detected in the mCherry (liposomes) and FITC (neurons) channels using a Beckman Coulter Gallios cytometer.

### *In vivo* biodistribution

4T1 cells were injected to the mammary fat pad of BALB/c female mice as was described above. Once the tumor reached 200 mm^3^, 200 µl of Rhodamine labeled liposomes were intravenously injected. 24 hours after injection mice were euthanized and tumors, livers, spleens, lungs, kidneys and hearts were extracted and imaged by *In-Vivo* Imaging System (IVIS). Moreover, liposomes accumulation was visualized using fluorescence histology imaging.

### IVIS imaging

As part of the biodistribution experiment, mice were euthanized and the extracted tissues were imaged *ex-vivo* using the IVIS SpectrumCT Pre-Clinical *In-Vivo* Imaging System (PerkinElmer, MA, USA) at an excitation of 570 nm and emission of 620 nm, binning 2, f-stop 2 and 8 seconds exposure (Fig. S4A). Quantitative data from all images was obtained using ROI tool in LivingImage software. A control (non-injected) mouse was used for analysis, and its average radiance of each tissue was subtracted from the average radiance of the injected mice.

In addition, at the end of the efficacy experiment 4T1 mCherry tumors were imaged by a whole-animal IVIS imaging at an excitation of 570 nm and emission of 620 nm, binning 8, f-stop 2 and 0.5 seconds exposure.

### Immunohistochemistry analysis

4T1 tumors were fixated using formalin solution, neutral buffered, 10% histological tissue fixative (Sigma-Aldrich, Rehovot, Israel) for at least 24 hour before embedding in paraffin and sectioned. Tissue sections were deparaffinized in a xylene ethanol gradient: xylene, xylene/ethanol (1:1 v/v), absolute ethanol, 95% ethanol, 70% ethanol, and 50% ethanol for 3 min each, and finally placed in distilled water (DW). Antigen retrieval was done in 10 mM tri-sodium citrate solution at pH 6 tittered with HCl. 2.5% ready-to-use normal goat serum (Vector Laboratories) was used for blocking. Incubation with primary antibody was at 4°C overnight with either rabbit anti-β III Tubulin or rabbit anti-PGP9.5 (1:2000, Abcam, Cambridge, UK). Tissue sections were rinsed 3 times with DW, and then were incubated for 30 minutes in 0.3% hydrogen peroxide solution for blocking of endogenous peroxidase activity Tissue sections were then washed in DW, and incubated with ready-to-use secondary goat anti-rabbit antibody conjugated to HRP (MP7451 kit, Vector Laboratories) for 40 minutes at RT. Tissue sections were rinsed 3 times with DW, and then for color development were incubated with DAB solution (SK4105 kit, Vector Laboratories) for 3 minutes, washed with DW, and counter stained with hematoxylin. Slides were scanned using 3DHistech Panoramic 250 Flash III automated slide scanner (3DHistech, Budapest, Hungary).

### Fluorescence immunohistochemistry analysis

4T1 tissue sections (10 µm thick) were fixed in 4% paraformaldehyde for 10 minutes, followed by 10 minutes of permeabilization with 0.1% Triton X-100. 10% normal donkey serum (Abcam, Cambridge, UK) in PBST (PBS with 0.025% Triton) was used for blocking for 2 hours. Tissue sections were incubated over night at 4°C in blocking buffer containing rabbit anti-β III Tubulin (1:1000, Abcam, Cambridge, UK) for neurons detection, and goat anti-CD31 (1:200, R&D systems, Minneapolis) for blood vessels detection. Following overnight incubation, sections were rinsed 3 times with PBST, then incubated with blocking buffer containing species-appropriate secondary antibodies; donkey anti-rabbit AlexaFluor 488 and donkey anti-goat AlexaFluor 647 (1:500, Abcam, Cambridge, UK) for 2 hours in RT. Tissue sections were rinsed 3 times with PBST, then mounted with DAPI-containing fluoromount (BioLegend, California), cover-slipped, and stored in the dark at 4°C until confocal imaging.

### *In vivo* therapeutic efficacy

4T1 mCherry cells injected subcutaneously as was described above. Once the tumor reached 100-200 mm^3^ (about 17 days post 4T1 injection), mice were divided to 4 different groups (5 mice each) as follows: Untreated, liposomal doxorubicin (L-DOX) (4 mg/kg-body-weight), and liposomal bupivacaine (L-BUP) in high (10 mg/kg) and low (3.5 mg/kg) dose. All of the treatments were administered by intravenous injection for 18 days. The high dose of L-BUP was injected twice a week, and the rest of the treatments were injected once a week. Mice weight and the tumor dimension were measured every two to three days. The tumor volume was measured using a caliper and calculated as (width^2^×length)/2. In order to compare mice weight and tumor dimension between the various treatments group, each mouse’ tumor size and the body weight was normalized to the initial size and weight that was measured at day 0 (the day before the first treatment).

### Metastases detection

Mice were euthanized, and the brain, lungs, bone marrow and tumor were extracted and held in RPMI at 4 °C until initiating the organ’s dissociation. Lungs and tumors were enzymatically and physically dissociated using a GentleMacs machine (Miltenyi Biotec, Bergisch Gladbach, Germany) and dissociation kit (Miltenyi Biotec) following the machine dissociation protocols. Brain tissue (*39, 40*) was cut to small pieces, and enzymatically digested by incubation in RPMI supplemented with 0.4 mg/mL collagenase D (Sigma-Aldrich), 0.2 mg/mL DNase I (Sigma-Aldrich) for 30 minutes in 37 °C at 100 rpm. Then the tissue was physically dissociated using the GentleMacs. The suspension was passed through a 70 μm cell strainer (BD Biosciences, CA, USA) and centrifuged for 400g for 5 minutes. The pellet was re-suspended with 7ml RPMI, then 3ml of 10:90 v/v of PBSX10:Percoll (GE Healthcare Bio-Sciences; Sigma-Aldrich) was added and the tube was gently mixed. The mixture was centrifuged for 400g for 25 minutes in 4 °C to form Percoll gradient, the supernatant was discarded and the pellet of the cells was re-suspended in PBS.

Bone marrow cells were harvested from the intact tibia and femur. The bone marrow was flushed out with PBS through a 25-gauge needle. Then, the PBS with the stromal cells were filtered in a 70 nm strainer and centrifuged at 500xg for 5 minutes.

The single-cell suspension of each tissue was obtained by passing the cell suspension through a 35 μm cell strainer (BD Biosciences, CA, USA). After dissociating of the organs into a single-cell suspension, the 4T1 mCherry cells were detected in the mCherry channel using FACS Aria and analyzed with FCS Express. Following acquisition of 10,000 single cells per sample, the average percentage of positive mCherry cells for each tissue was normalized to the average percentage measured at the control group.

### Quantification of immune cells in the tumor tissue

After dissociation of the tumor tissue into single cell suspension as was described above, cells suspension was incubated with CD45-FITC and CD3-APC diluted 1:200 for 1 hour in the dark on ice. Cells were washed three time with PBS, and analyzed using FACS Aria. For each test sample, 10,000 single cells were acquired, and data analysis was performed using FCS Express. The average percentage of the positive population was normalized to the average percentage measured at the control group

### Nerve detection

4T1 cells were injected to the mammary fat pad of BALB/c female mice as was described above. Once the tumor reached 300-500 mm^3^ (about 3 weeks from 4T1 injection), or the treatments of the efficacy experiment were completed (about 36 days from 4T1 injection) mice were euthanized. Single-cell suspension in addition to tissue sections were produced from the tumors, and nerve detection was conducted by immunohistochemistry and flow cytometry analyzes. As for the flow cytometry analysis, cells suspension was incubated with rabbit anti-Tyrosine hydroxylase (Abcam, Cambridge, UK), following a staining with donkey anti-rabbit Alexa Fluor 488 (Abcam, Cambridge, UK). Antibodies were diluted by 1:1000 in a blocking serum. The analysis was performed using BD LSR-II Analyzer (Biosciences, San Jose, CA, USA), and data analysis was obtained using FCS Express

### Statistical analysis

All statistical analysis: student’s t-test, one-way ANOVA and two-way ANOVA and three-way ANOVA were performed using Prism GraphPad software. Difference above *p*=0.05 considered statistically significant.

## Notes

### Competing Interest Statement

The authors have declared no competing interest.

