## Supplementary Information for "Targeting neurons in the tumor microenvironment with bupivacaine nanoparticles reduces breast cancer progression and metastases"

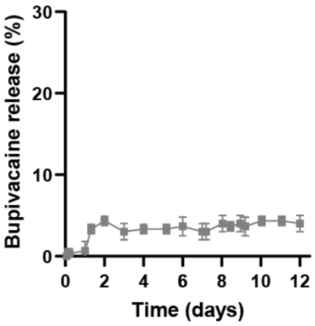


**Figure S1. Bupivacaine release profile in 4°C.** Liposomal bupivacaine release profile was conducted at 4°C for 12 days. Liposomes were composed of HSPC, cholesterol and DSPE-PEG2000 (molar ratio of 55:40:5). Results (three independent repetitions preformed in 3 replicates) are presented as mean±SD.

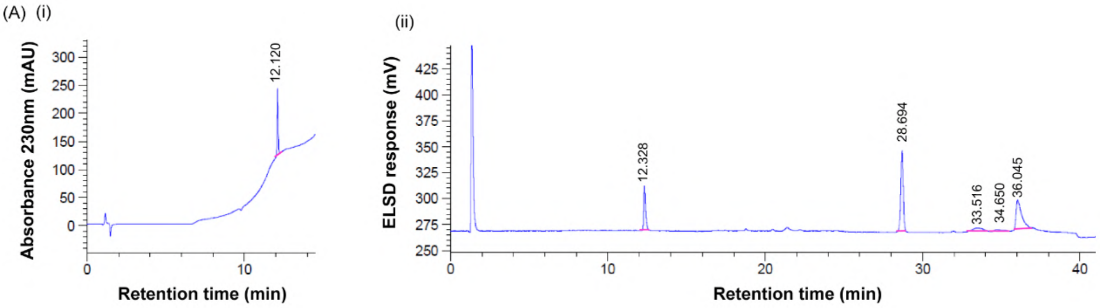


(B)

| | Bupivacaine | Cholesterol | DSPE-PEG2000 | HSPC | |
|---|---|---|---|---|---|
| | | | | 16:0 | 18:0 |
| **Retention time** (min) | 12.120 | 28.694 | 33.516 | 34.65 | 36.045 |
| **Area** (mAUxs or mVxs) | 755.851 | 874.181 | 205.467 | 922.275 | |
| **Concentration** (mg/ml) | 3.077 | 5.015 | 4.784 | 12.078 | |

**Figure S2. HPLC analysis of liposomal bupivacaine.** Chromatogram of liposomal bupivacaine using UV detector (A, i) and ELSD detector (A, ii). Analysis of the data regarding bupivacaine and the lipids obtained from UV and ELSD detectors respectively (B).

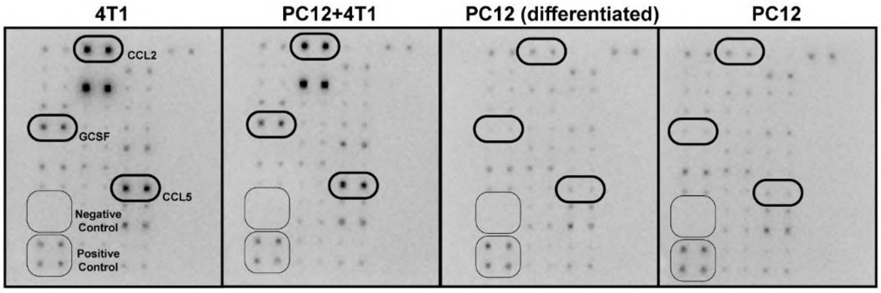


**Figure S3. Cancer cells stimulate neurite outgrowth through cytokines secretion.** Cancer cells secrete cytokines that promote neurites outgrowth, as was identified by cytokine antibody array. The analysis was conducted using conditioned media of different treatments after 72 hours from seeding.

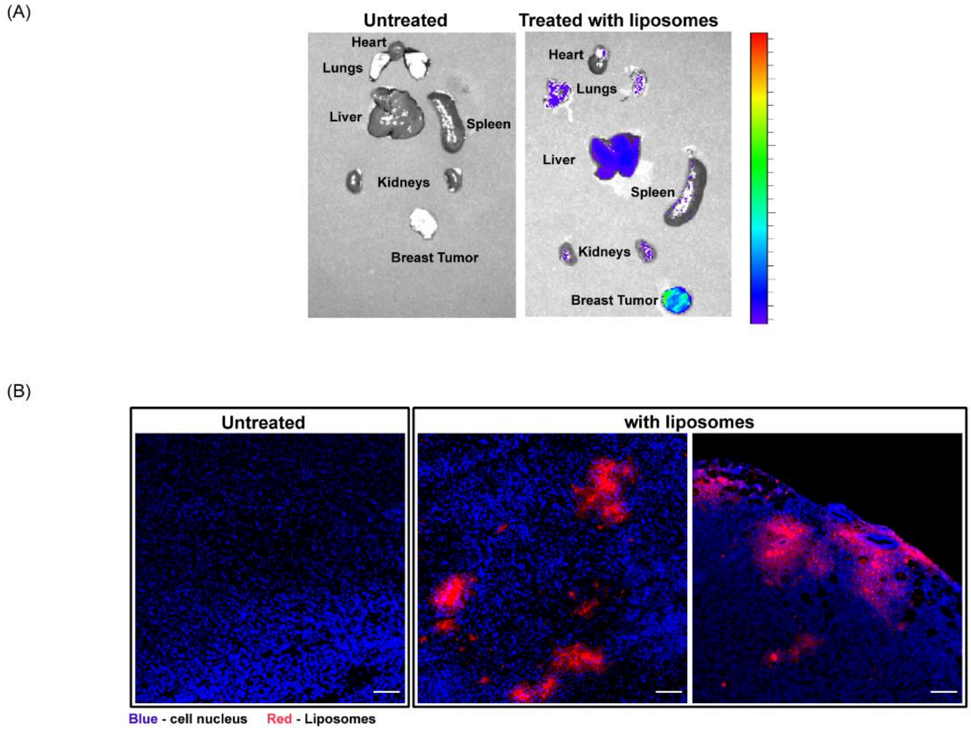


**Figure S4. Liposomal bupivacaine treatment inhibits tumor growth and metastases.** Liposomes labeled with Rhodamine were i.v. injected to mice bearing orthotopic 4T1 tumors, and after 24 hours their accumulation in different tissues was quantified using IVIS ex-vivo imaging (A). Liposome accumulation in the tumor tissue is also demonstrated by fluorescent histology images of the tumor (B, scale bar is 5cm).

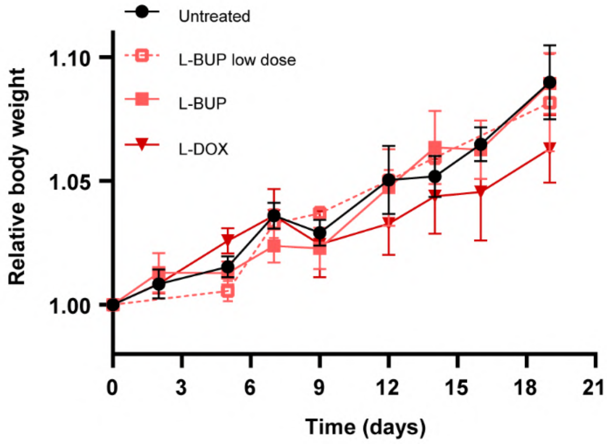


**Figure S5. Liposomal bupivacaine treatment does not affect mice's body weight.** No toxic effects were observed due to the treatments, and all mice's weight gradually increased. Body weight of mice was normalized to the initial weight at day 0. Results ($4 \le n \le 6$) are presented as mean±SEM.
